# Engagement in moderate-intensity physical activity supports overnight emotional memory retention in older adults

**DOI:** 10.1101/2024.08.16.608272

**Authors:** Miranda G. Chappel-Farley, Destiny E. Berisha, Abhishek Dave, Rachel M. Sanders, Christopher E. Kline, John C. Janecek, Negin Sattari, Kitty K. Lui, Ivy Y. Chen, Ariel B. Neikrug, Ruth M. Benca, Michael A. Yassa, Bryce A. Mander

**Author notes:** **Corresponding Author:** Miranda G. Chappel-Farley, Ph.D |.

## Abstract

**Importance:** Preserving the ability to vividly recall emotionally rich experiences contributes to quality of life in older adulthood. While prior work suggests that moderate-intensity physical activity (MPA) may bolster memory, it is unclear whether this extends to emotionally salient memories consolidated during sleep.

**Objective:** To investigate associations between engagement in physical activity (PA) and overnight emotional memory retention and examine whether theoretically replacing 30-minutes of lower-intensity activity with MPA is associated with better consolidation.

**Design, Setting, and Participants:** A cross-sectional study of 40 community-dwelling older adults free of neurological and psychiatric disorders. Data were collected from May 2018 to July 2022 and analyzed from January to July 2024.

**Exposures:** Participants completed an overnight polysomnography (PSG) with emotional memory tested before and after sleep and a self-report questionnaire assessing habitual PA.

**Main Outcome(s) and Measures(s):** Emotional memory performance was assessed via recognition memory or mnemonic discrimination performance. Overnight memory retention was calculated by subtracting immediate test from delayed test performance for both recognition memory and mnemonic discrimination, with more negative scores indicating lower memory retention. Frequency and duration of MPA, light-intensity PA, non-exertive activity, and sedentary behavior were calculated from the Community Health Activities Model Program for Seniors (CHAMPS) Activities Questionnaire for Older Adults. Isotemporal substitution modelling evaluated whether statistically reallocating time spent in sedentary and lower-intensity activity to MPA was associated with better overnight memory retention.

**Results:** Data from 40 participants were analyzed (□_age_=72.3±5.8, 26 female). Better overnight emotional recognition memory retention was associated with the frequency (β=0.663, SE=0.212, p=0.003) and duration (β=0.214, SE=0.101, p=0.042) of MPA. No relationships were found with mnemonic discrimination or neutral recognition memory. Statistically modelling the replacement of 30 minutes of lower-intensity activity with MPA was associated with better overnight retention of emotional memories (β=0.108, SE=0.048, p=0.030), but not neutral (β=-0.029, SE=0.069, p=0.679).

**Conclusions and Relevance:** MPA may enhance sleep-dependent consolidation of emotional memories in older adults. Modest increases in MPA may yield significant benefits for sleep-dependent emotional memory retention. These findings may guide interventions to preserve memory function and inform public health recommendations by demonstrating that substituting even short durations of low-intensity activity for MPA could produce significant cognitive gains relevant for maintaining quality of life in older adulthood.

## Introduction

As the global population continues to age,^1^ identifying protective measures to preserve quality of life and stave off memory decline has become of critical importance. Excessive sedentary behavior^2^ is highly prevalent among the elderly population, with some studies estimating that older adults are sedentary for 65-85% of the waking day.^3^ Sedentary behaviors contribute to significant physical and cognitive health risks, including increased mortality,^4^ impaired memory processing,^5^ and affective dysfunction and disorders.^6^ Aging is also associated with sleep changes, with more fragmented sleep and less time being spent in deeper sleep stages.^7^ Sedentary behavior may partially contribute to these changes, as increased sedentary behavior is linked to sleep efficiency.^8^ These behavioral shifts are particularly concerning given the critical role of sleep in the consolidation of memories.^9^ Disruption in sleep due to aging can impair overnight memory consolidation, contributing to memory deficits.^7^

Frequent engagement in physical activity, on the other hand, seems to play a protective role against age-related memory decline^10^ and may improve mood and affect.^11^ Indeed, research from both animal models and human participants provides support for brain and cognitive benefits of PA^12–16^, particularly in the domains of episodic memory^17^ and executive function.^18^ In fact, PA improves subjective sleep quality,^19^ sleep efficiency,^18^ and bolsters neural oscillations involved in sleep-dependent memory consolidation.^20^ However, not all memories are created equal, and physical activity^21^ and sleep may interactively contribute to the preferential consolidation and integrity of specific memory traces.

Research shows that memories containing emotional content are better remembered than non-emotional ones.^22^ Sleep appears to selectively benefit the retention of emotional memories,^23–25^ but there is age-related variability in this effect.^26^ Some work suggests that positive memories may not always exhibit a benefit from sleep^27^ and that older adults may actually better remember negative information.^28^ Traditional emotional memory recognition tests, however, do not assess the quality of memory representations. This can be measured via mnemonic discrimination—a behavioral proxy for hippocampal pattern separation which facilitates the storage of similar experiences as unique, high fidelity memory traces.^29^ While recognition memory for neutral events is generally preserved in aging,^30^ older adults exhibit worse emotional versus neutral memory recognition after a 24-hr consolidation period (which naturally encompasses sleep) and no consolidation benefit for emotional mnemonic discrimination.^31^ It is possible that physical activity may bolster consolidation for emotional recognition memory and mnemonic discrimination in older adults, as work in young adults suggests that PA and fitness support mnemonic discrimination immediately following encoding.^32–36^ Understanding these nuanced relationships among physical activity- and sleep-related consolidation of emotional memory representations is important, as maintaining high fidelity, vivid representations of experiences embedded in rich emotional context may provide adaptive value and enhance quality of life in older adulthood.^37,38^

Given that moderate-to-vigorous PA is consistently tied to better cognitive and health outcomes,^39^ an outstanding question is whether replacing sedentary behavior with moderate-intensity physical activity (MPA) could yield significant improvements in sleep-related consolidation of emotional memories in older adults. Advances in statistical modelling enable the estimation of how health or behavior might change when substituting the time spent performing one type of activity with another.^40^ This approach, called Isotemporal Substitution Modelling (ISM), aids in the identification of which activity substitutions are likely to facilitate better outcomes. A burgeoning literature using this theoretical modelling demonstrates that reallocating time spent in sedentary or non-exertive behavior to MPA improves a wide range of health outcomes.^41,42^ A recent study found that replacing sedentary time for light-intensity PA also improves episodic memory function in middle- and older-aged Latinx adults.^43^ However, this method has not yet been applied to sleep-related emotional memory consolidation in older adults, which is critical, as impairments in sleep-related memory processing contribute to age-related memory deficits. It is unknown whether replacing time spent in lower-intensity activities with MPA supports the consolidation of memory for emotionally laden experiences in the elderly. Understanding these relationships is essential, as these findings can inform the development of targeted interventions and inform public health recommendations aimed at preserving memory in late life.

To address these knowledge gaps, the current study assessed memory performance on an emotional variant of the mnemonic discrimination task (MDT) prior to and following overnight sleep, and self-reported habitual PA engagement over the four weeks prior to an in-lab polysomnography (PSG) study in sample of cognitively unimpaired older adults. Two primary hypotheses were tested: (1) that more frequent and longer duration engagement in MPA would be specifically associated with better overnight retention of emotional memories when assessing both mnemonic discrimination and recognition memory, and that (2) replacing lower-intensity activities with MPA would yield a significant sleep-related emotional memory retention benefit.

## Methods

### Study Participant Details

Forty cognitively unimpaired older adults between the ages of 60-85 years were recruited from the Biomarker Exploration in Aging, Cognition, and Neurodegeneration (BEACoN) cohort for an in-lab overnight polysomnography (PSG) assessment (□_age_=72.3±5.8, 26 Female, Mini Mental State Exam scores >24, **Table 1**). Participants were free of neurological and psychiatric disorders and were not taking sleep or cognition-affecting medications such as sedative hypnotics, non-SSRI antidepressants, anxiolytics, or neuroleptics. Participants did not have any prior diagnosis of sleep disorders other than sleep-disordered breathing (SDB) prior to participating and those with a prior SDB diagnosis were not being treated with continuous positive airway pressure. The severity of SDB was evaluated using the apnea-hypopnea index (AHI) which was derived from the PSG assessment and included as a covariate in all statistical models. Participants had not traveled more than three time zones within the three weeks prior to participating. On the day of their sleep study, participants were instructed to not consume caffeine any time after 9:00 AM. All participants provided written informed consent according to procedures approved by the University of California, Irvine Institutional Review Board.

**Table 1.** Sample characteristics.

| **Sample Characteristics** | **N=40** |
| --- | --- |
| Age, Mean [SD] | 72.3 [5.8] |
| Sex, No. Female (%) | 26 (65) |
| Years of Education, Mean [SD] | 16.7 [2.2] |
| BMI, Mean [SD] | 24.7 [5.6] |
| AHI, Mean [SD] | 13.4 [17.4] |
| Race, No. |  |
| African American or Black | 1 |
| Asian | 10 |
| White | 2 |
| More than one race | 26 |
| Unknown/Not Reported | 1 |
| Ethnicity, No. |  |
| Hispanic | 2 |
| Non-Hispanic | 38 |
| Sedentary Activity |  |
| Frequency (bouts/wk), Mean [SD] | 18.7 [18.0] |
| Duration (hours/wk), Mean [SD] | 17.0 [5.9] |
| Non-Exertive Activity |  |
| Frequency (bouts/wk), Mean [SD] | 7.8 [4.1] |
| Duration (hours/wk), Mean [SD] | 16.6 [9.8] |
| Light-Intensity Physical Activity |  |
| Frequency (bouts/wk), Mean [SD] | 11.8 [5.0] |
| Duration (hours/wk), Mean [SD] | 9.8 [7.0] |
| Moderate-Intensity Physical Activity |  |
| Frequency (bouts/wk), Mean [SD] | 8.8 [6.9] |
| Duration (hours/wk), Mean [SD] | 9.6 [9.1] |

### Community Healthy Activities Model Program for Seniors (CHAMPS) Activities Questionnaire for Older Adults

The CHAMPS Activities Questionnaire for Older Adults is a 40-item self-report questionnaire assessing engagement in various age-appropriate physical and lifestyle activities of various intensities during a ‘typical week’ when considering the prior four weeks.^44^ CHAMPS, as published, quantifies the frequency (number of times per week) and duration (total hours per week) of each reported activity to derive measures of frequency and duration of all exercise-related activities and moderate-intensity exercise-related activities. Given evidence that moderate-to-vigorous PA yields the greatest memory benefit^17^ our analyses were initially *a priori* restricted to the frequency and duration of moderate-intensity exercise-related activities, as published. The frequency and duration of light-intensity PA was derived to explore the specificity of PA intensity effects. A strength of this questionnaire is the inclusion of other, non-exertive activities that reduce socially desirable responding by allowing older adults who are less physically active to report other meaningful activities.^44^ Although not calculated in the original publication, the inclusion of these less physically active items allows for the quantification of frequency and duration of non-exertive and sedentary behavior. These variables were also derived and examined to determine the specificity of intensity-related findings. See **Tables S1-S3** for questionnaire scoring and details on the categorization of items into PA intensities. For ISM, each activity type was categorized (MPA as published in CHAMPS), and total weekly durations of each activity type were converted to average daily minutes and then rescaled to 30-minute intervals for substitution (**Table S2 & S4**). For greater detail on which activity types were categorized as sedentary, non-exertive, light-intensity, and moderate-intensity, see **Table S3**.

### Emotional Mnemonic Discrimination Task

Participants performed the emotional version of the Mnemonic Discrimination Task (eMDT) prior to and following overnight sleep.^45^ The eMDT is an established framework to measure emotionally-modulated hippocampal-dependent memory.^45,46^ Participants performed an incidental encoding phase followed by a surprise memory test prior to sleep and a delayed memory test the following morning. During the incidental encoding phase, 180 emotionally salient images were presented on black background for 4 seconds. While viewing, participants rated each image on valence (i.e., positive, negative, or neutral) using three individually colored orby button switches (P.I. Engineering, Williamston, MI). A white fixation cross was displayed on a black background for 500ms between image presentations. Immediately following the encoding phase, participants underwent a surprise memory test (immediate test phase) wherein they were presented with another 120 emotionally salient images split evenly among images they had seen during encoding (targets, 30), entirely new images (foils, 30), and images similar to those they saw during encoding (lures, 60). Lures could be somewhat similar to targets (low similarity, 30) or highly similar to targets (high similarity, 30). Participants indicated via button press whether they had seen the image before (‘old’) or whether it was an entirely new image (‘new). Following overnight PSG, participants completed another surprise memory test (delayed test phase), during which they viewed and responded to another 120 images split evenly among emotionally valent targets, foils, and lures as outlined above. No targets or corresponding lures presented in the immediate test phase were shown during the delayed test phase.

Recognition memory was calculated using *d’*, defined as z(‘old’|Targets) – z(‘old’|Foils). Mnemonic discrimination performance (behavioral proxy of hippocampal pattern separation) was calculated using the Lure Discrimination Index (LDI), defined as p(‘new’|lures) – p(‘new’|Targets), which corrects for response bias.^31^ LDI was calculated collapsing across similarity bins for each valence condition at both immediate and delayed testing timepoints; *d’* was calculated for each valence at both timepoints. Overnight memory retention for mnemonic discrimination ability and recognition memory were measured via overnight change in LDI (LDI_delayed test_-LDI_immediate test_) and *d’* (*d’*_delayed test_-*d’*_immediate test_), respectively. Both LDI and *d’* were analyzed to assess memory processing under conditions of moderate-to-high levels of interference (as measured by LDI) or no interference (as indicated by *d’*).

Multiple regression analyses were initially *a priori* restricted to LDI or d’ calculated for negative stimuli for two reasons: (1) prior work has shown that negative elements of an experience are preferentially consolidated during sleep^24,27,47^ and that, irrespective of age, positive experiences do not broadly exhibit a sleep benefit,^27^ and (2) positive and negative images in the eMDT are not matched on arousal making it difficult to appropriately disentangle the effects of positive versus negative valence.^46^ To show specificity of the relationships between PA and the consolidation of negative emotional stimuli, exploratory analyses were conducted using d’ for neutral images.

### Statistical Analyses

All analyses were conducted in SPSS v27.0 (IBM SPSS Statistics, Inc., Chicago, IL) and figures were created in RStudio v2022.12.0+353. Assumptions of multiple linear regression were checked. Frequency of MPA and duration of light-intensity PA were logarithm transformed (i.e., log(1+x)) and the duration of MPA was cube-root transformed to meet assumptions of normality.

Multiple regression analyses were used to examine relationships between self-reported PA and overnight memory retention while adjusting for age, sex, and AHI as covariates. Analyses were *a priori* restricted to negative images, but secondary multiple regression analyses adjusting for covariates was performed with positive stimuli (**Table S5**), which revealed no significant associations between PA and overnight positive memory retention. While prior work has shown that obstructive sleep apnea (OSA) severity impacts emotional memory consolidation and sleep neurophysiology,^48,49^ AHI was not associated with overnight emotional memory retention in the current sample (r=0.060, p=0.713). Despite the lack of a significant association between AHI and overnight emotional memory retention, AHI was included as a covariate due to the complexity of OSA effects on neurobiology and behavior.^49^ Primary analyses focused on MPA frequency (total number of bouts per week) and duration (total number of hours per week), as prior work has demonstrated that moderate-to-vigorous PA is most consistently tied to cognitive performance.^17,50,51^ Follow-up multiple regression analyses were performed additionally adjusting for sleep duration (minutes). To examine potential effects of PA intensity, control analyses examined effects of light-intensity PA and non-exertive activity frequency and/or duration adjusting for covariates. Follow-up multiple regression analyses adjusting for the same covariates were performed to determine whether relationships between MPA and overnight emotional memory retention were driven by relationships with immediate or delayed test recognition memory (*d’*) performance. Similarly, follow-up multiple regression analyses were performed adjusting for sleep duration. False Discovery Rate (FDR) correction was used to adjust for multiple comparisons for models predicting overnight change in emotional recognition memory from MPA frequency (3 comparisons: MPA, LPA, non-exertive) and MPA duration (3 comparisons: MPA, LPA, non-exertive).

### Isotemporal Substitution Model

Isotemporal substitution analyses evaluate the effect of reallocating a set duration of time from one activity type to another, while keeping total time constant.^40^ ISM was performed to determine whether reallocating 30 minutes of sedentary behavior, non-exertive activity, or light-intensity PA to MPA would yield an overnight emotional memory consolidation benefit. A 30-minute time allocation was selected based on previous literature^40^ and public health relevance.^52^ The ISM constructed for the current analyses can be expressed as:

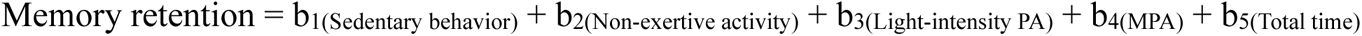

Where b_1_-b_5_ are the unstandardized regression coefficients. ISM is conducted by iteratively eliminating one activity component from the model and evaluating the remaining coefficients for each remaining activity type. The remaining coefficients represent the effect of substituting 30-minutes/day of the eliminated activity type for the remaining activities in the model, while holding total discretionary time constant. For example, by eliminating b_1_ (sedentary time) from the model, the remaining coefficients represent the effect of substituting 30 minutes of sedentary time for that respective activity (e.g., non-exertive activity, light-intensity PA, and MPA), while holding time spent in other activities and total time constant. Pearson’s correlations were performed to test for multicollinearity; importantly, average daily minutes spent in each activity type were not significantly correlated (See **Table S6**). Two separate ISMs were constructed separately to evaluate the effects of time reallocation on overnight emotional and neutral memory retention, respectively. Given that the intensity of some activities could not be determined given the lack of specificity into some aspects of the questionnaire (e.g., type of musical instrument played), we re-constructed ISM re-categorizing some activities up an intensity category to test for consistency in our findings (by MCF & CEK). Namely, we recategorized item 17 from sedentary to light-intensity and item 13 from non-exertive to light-intensity. These models produced similar results (see **Tables S7-S8**).

## Results

### Moderate-intensity physical activity is associated with emotional memory consolidation

First, we examined whether MPA frequency was associated with overnight retention of emotional mnemonic discrimination. A multiple regression model adjusting for age, sex and AHI (hereafter referred to as covariates) revealed no statistically significant association between MPA and overnight retention of emotional mnemonic discrimination, measured by overnight change in negative LDI (β=0.149, SE=0.101, p=0.149). Similarly, no relationship was found with overnight change in neutral LDI (β =0.025, SE=0.095, p=0.792). However, a regression model using overnight change in negative *d’* as the independent variable (IV) to measure recognition memory performance revealed that greater frequency of MPA was associated with better overnight emotional recognition memory retention (β=0.663, SE=0.212, p=0.003, FDR-adjusted p=0.0090; **Figure 1a**). This result remained statistically significant when sleep duration was added into the model as an additional covariate (β=0.666, SE=0.221, p=0.005). Importantly, this relationship was specific to negative emotional items, as MPA frequency was not associated with overnight change in neutral recognition memory *d’* (β=0.216, SE=0.308, p=0.487). This remained the case when sleep duration was added into the model (β =0.252, SE=0.321, p=0.438). The frequency of light-intensity PA (β=0.002, SE=0.018, p=0.891, FDR-adjusted p=0.891) and non-exertive activity (β=-0.020, SE=0.022, p=0.365, FDR-adjusted p=0.548) were not associated with overnight change in emotional recognition memory. When CHAMP items 13 and 17 were recategorized as light-intensity PA, the frequency of light intensity PA (β=0.005, SE=0.017, p=0.748) and non-exertive activity (β=-0.019, SE=0.022, p=0.399) still exhibited no statistically significantly associations with overnight retention of emotional recognition memory.

**Figure 1.**
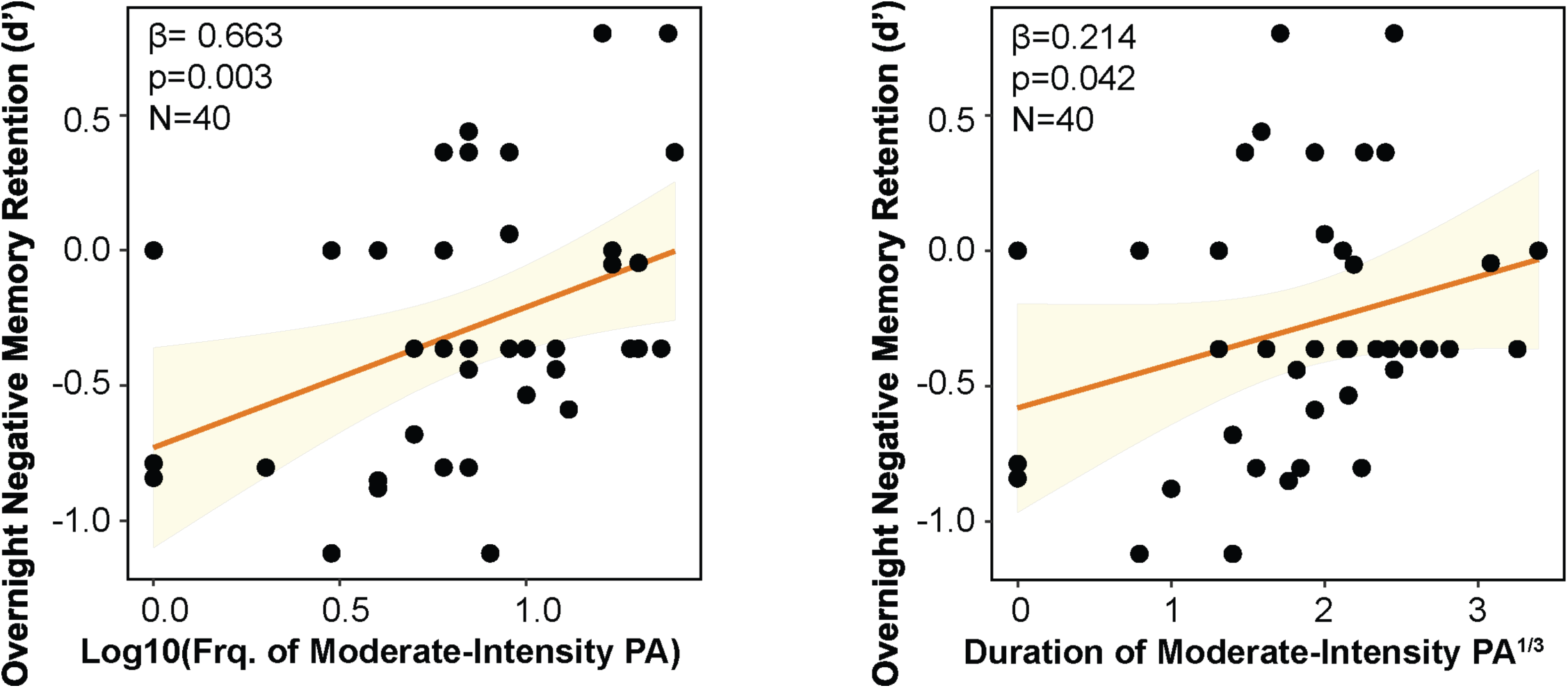
(A) More frequent and (B) longer duration moderate-intensity physical activity is associated with better overnight retention of emotional memories. Abbreviations: Frq—Frequency; PA—Physical activity

Next, the relationship between MPA duration and overnight retention of emotional recognition memory was tested. Greater duration of MPA was also associated with better overnight emotional recognition memory retention (*d’*), adjusting for covariates (β=0.214, SE=0.101, p=0.042; **Figure 1b**). However, this did not survive correction for multiple comparisons (FDR-adjusted p=0.126). This relationship was trending when sleep duration was added as an additional model covariate (β=0.210, SE=0.106, p=0.055). Of note, duration of light-intensity PA (β=0.006, SE=0.012, p=0.611, FDR-adjusted p=0.873) and non-exertive activity (β=-0.001, SE=0.009, p=0.873, FDR-adjusted p=0.873) were not associated with overnight emotional recognition memory retention. These results remained when CHAMP items 13 and 17 were recategorized as light-intensity PA (β=0.009, SE=0.012, p=0.471) and non-exertive activity (β=-0.001, SE=0.009, p=0.939). These results suggest that frequency and duration of MPA, specifically, supports overnight emotional recognition memory retention, but not mnemonic discrimination.

Follow-up multiple regression analyses were conducted to determine whether the relationship between MPA and overnight emotional recognition memory retention was driven by performance at immediate or delayed test. Neither the frequency (β=-0.242, SE=0.193, p=0.219) nor the duration (β=-0.029, SE=0.088, p=0.748) of MPA were associated with emotional recognition memory performance at the immediate testing timepoint, when adjusting for the same covariates. Results were similar when sleep duration was included as a covariate (MPA frequency: β=-0.225, SE=0.202, p=0.272; MPA duration: β=-0.016, SE=0.092, p=0.866). However, MPA frequency (β=0.421, SE=0.230, p=0.076) and duration (β=0.185, SE=0.103, p=0.082) exhibited trending relationships with performance at delayed test, further suggesting that MPA engagement supports the consolidation process in older adults. This finding was similar when sleep duration was included as an additional covariate in these models (MPA frequency: β=0.441, SE=0.240, p=0.075; MPA duration: β=0.195, SE=0.108, p=0.081).

To explore whether these relationships were due to MPA effects on sleep architecture, Pearson’s correlations were tested to examine if MPA frequency or duration were associated with measures of global sleep architecture. These models revealed no statistically significant associations between MPA frequency or duration with sleep duration, sleep efficiency (time spent asleep/time in bed), wake after sleep onset, percentage of time spent in non-rapid eye movement sleep (NREM) stage 1, stage 2, stage 3, or REM (**Tables S9-S10**).

### Replacing 30 minutes of lower-intensity activity with moderate-intensity physical activity yields an emotional memory consolidation benefit

ISM was performed to determine whether replacing 30 minutes of lower-intensity activity (i.e., sedentary behavior, non-exertive activity, light-intensity PA) with MPA would be associated with better negative emotional memory consolidation. Reallocating 30-minutes of non-exertive activity to MPA was associated with better overnight emotional memory retention (β=0.108, SE=0.048, p=0.030; **Table 2, Model B**). Reallocation of non-exertive activity to either sedentary time or light-intensity PA did not impact overnight emotional memory retention. Reallocation of 30 minutes of light-intensity PA to moderate-intensity activity showed a trending association with better overnight emotional memory retention (β=0.100, SE=0.059, p=0.099; **Table 2, Model C**). Importantly, the impact of time reallocation on overnight memory processing was specific to emotional events, as no relationships were observed with neutral stimuli (**Table 3**). When CHAMPS items 13 and 17 were recategorized as light-intensity PA, the ISM analyses produced similar findings (**Tables S7-S8**). These findings provide further support for the specificity of MPA effects on sleep-related consolidation of emotional memories and suggest that replacing even a short duration of lower-intensity activity for MPA enhances in emotional memory consolidation.

**Table 2.** Isotemporal substitution model: The effect of reallocating 30-min of time spent engaged in sedentary behavior, non-exertive activities, light-intensity physical activity, and moderate-intensity physical activity on overnight emotional memory retention (N=40).

| Isotemporal Substitution Model | Type of Activity |  |  |  |  |  |  |  |  |  |  |  |
| --- | --- | --- | --- | --- | --- | --- | --- | --- | --- | --- | --- | --- |
|  | SED |  |  | Non-Exertive |  |  | LPA |  |  | MPA |  |  |
| | $\beta$ | SE | p | $\beta$ | SE | p | $\beta$ | SE | p | $\beta$ | SE | p |
| Model A: Substitution of SED | — | — | — | -0.082 | 0.051 | 0.117 | -0.073 | 0.060 | 0.235 | 0.027 | 0.063 | 0.675 |
| Model B: Substitution of Non-Exertive | 0.082 | 0.051 | 0.117 | — | — | — | 0.009 | 0.050 | 0.862 | 0.108 | 0.048 | 0.030* |
| Model C: Substitution of LPA | 0.073 | 0.060 | 0.235 | -0.009 | 0.050 | 0.862 | — | — | — | 0.100 | 0.059 | 0.099<sup>t</sup> |
| Model D: Substitution of MPA | -0.027 | 0.063 | 0.675 | -0.108 | 0.048 | 0.030* | -0.100 | 0.059 | 0.099<sup>t</sup> | — | — | — |

Data are unadjusted for covariates. Negative values indicate worse overnight emotional memory retention. <sup>t</sup>p<0.10, \*p<0.05, \*\*p<0.01 Abbreviations: SED—sedentary behavior; LPA—light-intensity physical activity; MPA—moderate intensity physical activity; $\beta$ —unstandardized regression coefficient; SE—standard error; p—p-value

**Table 3.** Isotemporal substitution model: The effect of reallocating 30-min of time spent engaged in sedentary behavior, non-exertive activities, light-intensity physical activity, and moderate-intensity physical activity on overnight neutral memory retention (N=40).

| Isotemporal Substitution Model | Type of Activity |  |  |  |  |  |  |  |  |  |  |  |
| --- | --- | --- | --- | --- | --- | --- | --- | --- | --- | --- | --- | --- |
|  | SED |  |  | Non-Exertive |  |  | LPA |  |  | MPA |  |  |
| | $\beta$ | SE | p | $\beta$ | SE | p | $\beta$ | SE | p | $\beta$ | SE | p |
| Model A: Substitution of SED | — | — | — | 0.051 | 0.073 | 0.490 | 0.087 | 0.087 | 0.323 | 0.022 | 0.091 | 0.808 |
| Model B: Substitution of Non-Exertive | -0.051 | 0.073 | 0.490 | — | — | — | 0.036 | 0.506 | 0.616 | -0.029 | 0.069 | 0.679 |
| Model C: Substitution of LPA | -0.087 | 0.087 | 0.323 | -0.036 | 0.072 | 0.616 | — | — | — | -0.065 | 0.085 | 0.449 |
| Model D: Substitution of MPA | -0.022 | 0.091 | 0.808 | 0.029 | 0.069 | 0.679 | 0.065 | 0.085 | 0.449 | — | — | — |

Data are unadjusted for covariates. Negative values indicate worse overnight neutral memory retention. <sup>t</sup>p<0.10, \*p<0.05, \*\*p<0.01 Abbreviations: SED—sedentary behavior; LPA—light-intensity physical activity; MPA—moderate intensity physical activity; $\beta$ —unstandardized regression coefficient; SE—standard error; p—p-value

## Discussion

Results from the current study suggest that higher engagement in MPA (both frequency and duration) supports sleep-related emotional memory consolidation in older adults. Multiple regression analyses revealed that greater MPA was associated with better overnight emotional memory retention. ISM demonstrated that replacing 30-minutes of non-exertive activity with MPA yields an improvement in overnight emotional memory consolidation. Notably, each of these findings were specific to memories containing emotional content. These findings provide the first evidence that reallocating just 30-minutes of time spent in non-exertive activities to MPA yields an emotional memory consolidation benefit in older adults, a population at risk for cognitive decline. Targeted interventions and policies aimed at promoting memory preservation in older adults should include the substitution of more sedentary behaviors with MPA, even for short durations.

Prior work in older adults using ISM has shown that substituting 30-minutes of lower-intensity activity (i.e., sedentary behavior, light-intensity physical activity) with MPA is associated with better mental health,^53^ executive function,^43^ self-regulation,^54^ and episodic memory.^43^ The current findings build on this literature, showing that substituting just 30-minutes of non-active time with MPA may result in significant improvements in sleep-dependent consolidation of emotional memories in older adults. Notably, CHAMPS MPA items of the CHAMPS questionnaire constitute aerobic PA, which has been consistently tied to better memory function. In fact, several studies even suggest that aerobic PA may bias offline processing towards retention of emotional stimuli via changes in stress hormone action, neurotransmitter release, increased levels of brain-derived neurotrophic factor (BDNF), and changes in autonomic activity.^21,55,56^ PA appears to have time-dependent or cumulative effects on memory processing depending on the frequency of engagement.^57^ Acute exercise seems to yield the greatest memory boost when it is temporally coupled to learning, and recent work has demonstrated that a post-encoding aerobic exercise bout selectively promotes the consolidation of emotional experiences.^56^ In contrast, the effects of long-term PA appear to be cumulative and less time-dependent, driving changes in the neurophysiology of the medial temporal lobe (MTL) memory system over time with repeated bouts facilitating maintenance of memory enhancement.^57,58^ In line with this notion, our findings that MPA engagement (frequency and duration) is significantly associated with overnight emotional recognition memory consolidation and delayed retrieval—and not performance at immediate test—suggest that PA supports consolidation during sleep rather than simply enhancing encoding processes in older adulthood.

This study found that PA was specifically associated with emotional recognition memory and not emotional mnemonic discrimination ability. Furthermore, findings suggest that while MPA supports sleep-related memory consolidation, sedentary behavior and light-intensity activity do not necessarily actively impair this process, as evidenced by a lack of negative correlations between these measures. These findings underscore that engagement in MPA may bolster consolidation processes occurring during sleep in older adults. The finding that MPA was not associated with mnemonic discrimination performance contrasts research conducted in young adults, though these studies did not assess mnemonic discrimination performance over a period of sleep.^32–35^ While a recent study in physically active older adults found that object recognition memory significantly improved following a moderate-intensity aerobic exercise bout, mnemonic discrimination did not.^59^ However, findings in this domain are generally mixed.^60^ One study in initially inactive older adults found improved mnemonic discrimination performance following a chronic high-intensity exercise paradigm—an effect related to improved cardiorespiratory fitness^61^—while other studies in physically fit older adults found no relationship with cardiorespiratory fitness or any intervention-related changes in performance.^62–64^ On the other hand, another study found improved mnemonic discrimination after one bout of light-intensity aerobic exercise; however, the fitness level of participants prior to participation was not assessed.^65^ It is important to note that participants in the current study were relatively physically active, with 80% of the study participants meeting guidelines for MPA ( ≥150 min per week).^66^ While several of these prior studies did not directly compare recognition memory and mnemonic discrimination directly,^61–64^ our findings are consistent with those including samples of physically active older adults in that habitual PA was not associated with mnemonic discrimination performance.

The potential dissociation between mnemonic discrimination and recognition memory in our sample, and other works,^59^ is interesting and may highlight the importance of considering the baseline fitness level of participants. Perhaps, in physically active older adults, PA-related memory improvements extend beyond the hippocampus (which facilitates mnemonic discrimination) to other MTL structures to drive improvements in other facets of memory processing that support recognition memory. One possibility is that MPA bolsters familiarity, which is a key component of recognition memory that is less reliant on the hippocampus and more closely tied to perirhinal and parahippocampal cortex function.^67,68^ Indeed, MPA has been shown to strengthen functional connectivity in perirhinal and parahippocampal cortices,^17^ and PA increases parahippocampal BOLD-signal activation, angiogenesis, and gray matter volumes.^21^ Additionally, given that older adults often rely on familiarity-based strategies to compensate for memory deficits,^69^ it is possible that engagement in MPA may bolster this adaptive function. Future studies should investigate whether acute manipulation of physical activity bolsters sleep-dependent emotional memory consolidation

This study is the first to demonstrate that MPA specifically supports sleep-related emotional memory retention. The use of the emotional mnemonic discrimination task allowed us to investigate valence-specific differences, as well as differential effects on recognition memory and mnemonic discrimination. The use of the CHAMPS questionnaire enabled us to probe different aspects of PA across the human movement spectrum. The use of ISM in this context provided novel insights into the effects of reallocating sedentary time to physical activity, demonstrating that replacing just 30-minutes of low-intensity activity for moderate-intensity physical activity yields an emotional memory consolidation benefit in older adults. However, this study is not without limitations. PA was assessed using a self-report questionnaire; while allowing for the quantification of self-reported frequency and duration of engagement, this limited our ability to differentiate different activity types due to their inclusion in a single questionnaire item and limited our ability to evaluate the effects of physical fitness. Furthermore, as the questionnaire assessed typical activity engagement over the past four weeks, PA engagement on the day of the overnight sleep study visit was not assessed, thereby limiting the ability to differentiate acute versus chronic effects on memory performance. Additionally, total time spent in each activity was assumed to be equal across each day of the week for ISM. Future work should utilize device-based monitoring to assess daily sedentary behavior and PA engagement, as well as objective fitness assessments. Participants in the current study were relatively active, as evidenced by meeting PA guidelines for moderate intensity activity, and exhibited SDB symptoms, both of which may independently or interactively influence neurophysiology associated with sleep-dependent memory processing. For this reason, AHI was included as a covariate in our analyses. While the average AHI in our sample was representative of community dwelling older adults,^70^ future studies should examine whether these relationships differ in older adult populations with lower SDB burden. While global measures of sleep architecture were not associated with MPA engagement, future studies utilizing larger samples should investigate whether MPA is associated with the expression of sleep oscillations implicated in the consolidation process, such as slow oscillations and sleep spindles.^9^ Lastly, the cross-sectional design limits our ability to definitively infer causal relationships between PA and emotional memory consolidation. Future studies should use objective measures of fitness and PA (i.e., VO_2_ max, actigraphy) and/or exercise manipulations in a larger, more diverse sample free of SDB to determine whether reallocating 30-minutes of low intensity activity improves memory function. Despite these limitations, the findings from the current study have important public health implications and can inform interventions and recommendations to preserve memory function in later life.

## Funding Information and Acknowledgements

This research was supported by grants NIA F31AG074703, NIA F31AG084308, NHLBI T32HL082610, NIA R21AG07955, NIA R01AG053555, NIA K01AG058353, and the American Academy of Sleep Medicine Strategic Research Award. We would like to thank all of the research participants who generously devoted their time to this study, as well as the laboratory technicians, research specialists, and undergraduate research assistants to aided in data collection.

