## Supplemental Material for "Engagement in moderate-intensity physical activity supports overnight emotional memory retention in older adults"

**Table S1.** CHAMPS outcome measures and scoring criteria.

| Outcome Variable | Item Number | Coding |
|---|---|---|
| *Exercise-related Activities* | | |
| Frequency of all moderate-intensity exercise-related activities[1] | 7, 9, 14-16, 19, 21, 23-26, 29-33, 36- 38, 40 | Sum frequency scores/week for each for the subset of activities categorized as moderate intensity (MET >= 3.0) |
| Duration of all moderate-intensity exercise-related activities[1] | 7, 9, 14-16, 19, 21, 23-26, 29-33, 36- 38, 40 | Sum the numeric duration variable for the subset of activities categorized as moderate intensity (MET >= 3.0) |
| Frequency of light-intensity exercise related activities | 10, 20, 22, 27, 28, 34, 35, 39 | Sum frequency scores/week for each for the subset of activities categorized as light intensity (MET < 3.0) |
| Duration of all light-intensity exercise-related activities | 10, 20, 22, 27, 28, 34, 35, 39 | Sum the numeric duration variable for the subset of activities categorized as light intensity (MET < 3.0) |
| *Non-Exertive Activities* | | |
| Frequency of non-exertive behavior | 1-5, 11-13 | Sum frequency scores/week for each activity categorized as non-exertive (allow those with missing data to be included in the sum) |
| Duration of non-exertive behavior | 1-5, 11-13 | Sum the numeric duration variable for the subset of activities categorized as non-exertive |
| *Sedentary Behavior* | | |
| Frequency of Sedentary Behavior | 6, 8, 17, 18 | Sum frequency scores/week for each activity categorized as sedentary (allow those with missing data to be included in the sum) |
| Duration of Sedentary Behavior | 6, 8, 17, 18 | Sum the numeric duration variable for the subset of activities categorized as sedentary |

**Table S2.** Calculation of outcome variables from CHAMPS for ISM.

| Outcome Variable | Item Numbers | Coding |
|---|---|---|
| Duration of Sedentary Behavior | 6, 8, 17, 18 | Sum the numeric duration variable for the subset of activities noted as sedentary. Divide this numeric by 7 to calculate average hours/day. Convert to minutes/day. Divide by 30 to rescale to 30-minute intervals for substitution. |
| Duration of Non-Exertive Behavior | 1, 2, 3, 4, 5, 11, 12, 13 | Sum the numeric duration variable for the subset of activities noted as non-exertive. Divide this numeric by 7 to calculate average hours/day. Convert to minutes/day. Divide by 30 to rescale to 30-minute intervals for substitution. |
| Duration of Light-Intensity Physical Activity | 10, 20, 22, 27, 28, 34, 35, 39 | Sum the numeric duration variable for the subset of activities noted as light intensity (MET < 3.0). Divide this numeric by 7 to calculate average hours/day. Convert to minutes/day. Divide by 30 to rescale to 30-minute intervals for substitution. |
| Duration of Moderate-Intensity Physical Activity | 7, 9, 14, 15, 16, 19, 21, 23, 24, 25, 26, 29, 30, 31, 32, 33, 36, 37, 38, 40 | Sum the numeric duration variable for the subset of activities noted as moderate intensity (MET >= 3.0). Divide this numeric by 7 to calculate average hours/day. Convert to minutes/day. Divide by 30 to rescale to 30-minute intervals for substitution. |

**Table S3.** Categorization of CHAMPS Questionnaire items

| Sedentary Behaviors | Non-Exertive Activities | Light-Intensity Activities | Moderate Intensity Activities* |
|---|---|---|---|
| 6. Using a computer<br>8. Woodworking/needlework/drawing/arts and crafts<br>17. Play a musical instrument<br>18. Reading | 1. Visit friends and family<br>2. Go to the senior center<br>3. Volunteer work<br>4. Attend church<br>5. Attend club or group meetings<br>11. Attend a concert, movie, lecture, or sport event<br>12. Play cards, bingo, or board games with other people<br>Shoot pool or billiards | 10. Play golf, riding a cart<br>20. Light housework<br>22. Light gardening<br>27. Walk to do errands<br>28. Walk leisurely for exercise or pleasure<br>34. Stretching or flexibility exercises<br>35. Yoga or Tai-chi<br>39. General conditioning exercises, such as light calisthenics or chair exercises | 7. Dance<br>9. Play golf, carrying or pulling equipment<br>14. Play singles tennis<br>15. Play doubles tennis<br>16. Ice/roller/in-line skate<br>19. Do heavy housework<br>21. Heavy gardening (spading or raking)<br>23. Work on car, truck, lawn mower, other machinery<br>24. Jog or run<br>25. Walk or hike uphill<br>26. Walk fast or briskly for exercise<br>29. Ride a bike<br>30. Aerobics including rowing or step machines<br>31. Water exercises<br>33. Swim gently<br>32. Swim moderately or fast<br>36. Aerobics or aerobic dancing<br>37. Moderate-to heavy strength training<br>38. Light strength training<br>40. Basketball, soccer, or racquetball |

*As published

**Table S4.** Calculation of CHAMPS variables for ISM with recategorization of items 13 and 17 to light-intensity activity.

| Outcome Variable | Item Numbers | Coding |
|---|---|---|
| Duration of Sedentary Behavior | 6, 8, 18 | Sum the numeric duration variable for the subset of activities noted as sedentary. Divide this numeric by 7 to calculate average hours/day. Convert to minutes/day. Divide by 30 to rescale to 30-minute intervals for substitution. |
| Duration of Non-Exertive Behavior | 1, 2, 3, 4, 5, 11, 12 | Sum the numeric duration variable for the subset of activities noted as non-exertive. Divide this numeric by 7 to calculate average hours/day. Convert to minutes/day. Divide by 30 to rescale to 30-minute intervals for substitution. |
| Duration of Light-Intensity Physical Activity | 10, 13, 17, 20, 22, 27, 28, 34, 35, 39 | Sum the numeric duration variable for the subset of activities noted as light intensity (MET < 3.0). Divide this numeric by 7 to calculate average hours/day. Convert to minutes/day. Divide by 30 to rescale to 30-minute intervals for substitution. |
| Duration of Moderate-Intensity Physical Activity | 7, 9, 14, 15, 16, 19, 21, 23, 24, 25, 26, 29, 30, 31, 32, 33, 36, 37, 38, 40 | Sum the numeric duration variable for the subset of activities noted as moderate intensity (MET >= 3.0). Divide this numeric by 7 to calculate average hours/day. Convert to minutes/day. Divide by 30 to rescale to 30-minute intervals for substitution. |

**Table S5.** Results from multiple regression analyses predicting overnight change in positive recognition memory performance and overnight change in positive mnemonic discrimination performance from the frequency and duration of moderate-intensity physical activity, adjusting for age, sex, and the apnea-hypopnea index.

| | Overnight change in positive *d'* | Overnight change in positive LDI |
|---|---|---|
| Frequency of MPA | β=-0.304, SE=0.363, p=0.407 | β=-0.034, SE=0.092, p=0.714 |
| Duration of MPA | β=-0.173, SE=0.162, p=0.292 | β=-0.022, SE=0.041, p=0.599 |

Abbreviations: MPA—Moderate-intensity physical activity; LDI—Lure Discrimination Index; β—unstandardized regression coefficient; SE—standard error; p—p-value

**Table S6.** Pearson's correlation values between average daily minutes spent in each activity type.

| | Daily Minutes of Moderate-Intensity Physical Activity | Daily Minutes of Light-Intensity Physical Activity | Daily Minutes of Social Non-Exertive Activity | Daily Minutes of Sedentary Behavior |
|---|---|---|---|---|
| Daily Minutes of Moderate-Intensity Physical Activity | — | r=0.221, p=0.170 | r=0.109, p=0.501 | r=0.206, p=0.202 |
| Daily Minutes of Light-Intensity Physical Activity | — | — | r=0.185, p=0.253 | r=0.124, p=0.444 |
| Daily Minutes of Social Non-Exertive Activity | — | — | — | r=0.027, p=0.868 |
| Daily Minutes of Sedentary Behavior | — | — | — | — |

**Table S7.** Isotemporal substitution model with items 13 & 17 recategorized as light-intensity: The effect of reallocating 30-min of time spent engaged in sedentary behavior, non-exertive activities, light-intensity physical activity, and moderate-intensity physical activity on overnight emotional memory retention (N=40).

| Isotemporal Substitution Model | Type of Activity | | | | | | | | | | | |
|---|---|---|---|---|---|---|---|---|---|---|---|---|
| | SED | | | Non-Exertive | | | LPA | | | MPA | | |
| | β | SE | p | β | SE | p | β | SE | p | β | SE | p |
| Model A: Substitution of SED | — | — | — | -0.091 | 0.058 | 0.122 | -0.089 | 0.071 | 0.219 | 0.019 | 0.067 | 0.778 |
| Model B: Substitution of Non-Exertive | 0.091 | 0.057 | 0.122 | — | — | — | 0.002 | 0.053 | 0.976 | **0.110** | **0.048** | **0.029*** |
| Model C: Substitution of LPA | 0.089 | 0.071 | 0.219 | -0.002 | 0.053 | 0.976 | — | — | — | **0.108** | **0.061** | **0.083[t]** |
| Model D: Substitution of MLA | -0.019 | 0.067 | 0.778 | **-0.110** | **0.048** | **0.029*** | **-0.108** | **0.061** | **0.083[t]** | — | — | — |

Data are unadjusted for covariates. Negative values indicate worse overnight emotional memory retention. [t]$p<0.10$, *$p<0.05$, **$p<0.01$
Abbreviations: SED—sedentary behavior; LPA—light-intensity physical activity; MPA—moderate intensity physical activity; β—unstandardized regression coefficient; SE—standard error; p—p-value

**Table S8.** Isotemporal substitution model with items 13 & 17 recategorized as light-intensity: The effect of reallocating 30-min of time spent engaged in sedentary behavior, non-exertive activities, light-intensity physical activity, and moderate-intensity physical activity on overnight neutral memory retention (N=40).

| Isotemporal Substitution Model | Type of Activity | | | | | | | | | | | |
|---|---|---|---|---|---|---|---|---|---|---|---|---|
| | SED | | | Non-Exertive | | | LPA | | | MPA | | |
| | β | SE | p | β | SE | p | β | SE | p | β | SE | p |
| Model A: Substitution of SED | — | — | — | 0.030 | 0.084 | 0.726 | 0.032 | 0.104 | 0.757 | -0.006 | 0.098 | 0.956 |
| Model B: Substitution of Non-Exertive | -0.030 | 0.084 | 0.726 | — | — | — | 0.003 | 0.078 | 0.971 | -0.035 | 0.071 | 0.623 |
| Model C: Substitution of LPA | -0.032 | 0.104 | 0.757 | -0.003 | 0.078 | 0.971 | — | — | — | -0.038 | 0.089 | 0.672 |
| Model D: Substitution of MPA | 0.006 | 0.098 | 0.958 | 0.035 | 0.071 | 0.623 | 0.038 | 0.089 | 0.672 | — | — | — |

Data are unadjusted for covariates. Negative values indicate worse overnight neutral memory retention. $^{t}p<0.10$, $^{*}p<0.05$, $^{**}p<0.01$
Abbreviations: SED—sedentary behavior; LPA—light-intensity physical activity; MPA—moderate intensity physical activity; β—unstandardized regression coefficient; SE—standard error; p—p-value

**Table S9.** Pearson's correlation analyses predicting percentage of time spent in each sleep stage, sleep efficiency, and WASO from the frequency of moderate-intensity physical activity.

| | TST | NREM1 | NREM2 | NREM3 | REM | SE | WASO |
|---|---|---|---|---|---|---|---|
| MPA Frequency | r=-0.161, p=0.320 | r=0.051, p=0.755 | r=0.259, p=0.107 | r=-0.158, p=0.329 | r=-0.183, p=0.259 | r=-0.077, p=0.638 | r=0.114, p=0.484 |

Abbreviations: MPA—Moderate-intensity physical activity; TST— Total sleep time (minutes); NREM—Non-rapid eye movement sleep; REM—Rapid eye movement sleep; SE—Sleep Efficiency; WASO—Wake after sleep onset

**Table S10**. Pearson's correlation analyses predicting percentage of time spent in each sleep stage, sleep efficiency, and WASO from the duration of moderate-intensity physical activity.

| | TST | NREM1 | NREM2 | NREM3 | REM | SE | WASO |
|---|---|---|---|---|---|---|---|
| MPA Duration | r=-0.250, p=0.119 | r=0.128, p=0.432 | r=0.257, p=0.109 | r=-0.149,p=0.359 | r=-0.264,p=0.100 | r=-0.119, p=0.466 | r=0.198, p=0.222 |

Abbreviations: MPA—Moderate-intensity physical activity; TST— Total sleep time (minutes); NREM—Non-rapid eye movement sleep; REM—Rapid eye movement sleep; SE—Sleep Efficiency; WASO—Wake after sleep onset
